# Discovery of *Pandoraea pnomenusa* RB38 *N*-acyl homoserine Lactone Synthase (PpnI) and its Complete Genome Sequence Analysis

**DOI:** 10.1101/020180

**Authors:** Kok-Gan Chan, Robson Ee, Kah-Yan How, Siew-Kim Lee, Wai-Fong Yin, Yan-Lue Lim

## Abstract

In this study, we sequenced the genome of *P. pnomenusa* RB38 and reported the finding of a pair of cognate *luxI/R* homologs which we firstly coined as *ppnI*, which is found adjacent to a *luxR* homolog, *ppnR*. An additional orphan *luxR* homolog, *ppnR*2 was also discovered. Multiple sequence alignment revealed that PpnI is a distinct cluster of AHL synthase compared to those of its nearest phylogenetic neighbor, *Burkholderia* spp. When expressed heterologously and analysed using high resolution tandem mass spectrometry, PpnI directs the synthesis of *N*-octanoylhomoserine lactone (C8-HSL). To our knowledge, this is the first documentation of the *luxI/R* homologs of the genus of *Pandoraea*.

## INTRODUCTION

The theory of “quorum sensing” (QS) was coined in the late nineties describing bacterial cell-to-cell communication for various gene expression regulations (Bainton *et al.* 1992; Miller & Bassler 2001; Schauder & Bassler 2001). This communication is accomplished through secretion and detection of small hormone-like chemical molecules known as autoinducers which facilitate intra- and inter-species microbial communications. There are different classes of autoinducers where upon reaching a threshold concentration, these signaling molecules activate and stimulate a wide variety of gene expression (Davies *et al.* 1998; Williams *et al.* 2007). The most studied QS molecules is *N*-acyl homoserine lactone (AHL) which is secreted by Gram-negative proteobacteria especially in the class of alpha-, beta- and gamma-proteobacteria subdivisions. AHL typically consists of a homoserine lactone moiety (Williams *et al.* 2007) and an *N*-acyl side chain with various chain length (C4-C18), degree of saturation at C-3 position and presence of a hydroxy-, oxo- or no substituent at the C3 position (Chhabra *et al.* 2005). AHL synthase, also known as the LuxI homologs, together with the AHL receptor protein known as LuxR homologs, are two typical principal protein families in AHLs QS system. Briefly, in this QS system, AHLs are secreted by LuxI homologs until a threshold concentration of AHL is attained before they bind to LuxR homologs and subsequently activate a cascade of QS-regulated gene expression (Fuqua, Parsek & Greenberg 2001; Swift *et al.* 2001; Swift *et al.* 1996).

*Pandoraea* was believed to be originated from the term “Pandora box” which referred to the source of all evil in Greek mythology. Predominantly isolated from cystic fibrosis (CF) patients, *Pandoraea* species were also recovered from other clinical specimen and soil environment (Coenye *et al.* 2000; Daneshvar *et al.* 2001). Clinical manifestations of this terrorizing pathogen revolved around nosocomial infections with its capability to deteriorate lung function (Caraher *et al.* 2008; Costello *et al.* 2011; Stryjewski *et al.* 2003) and even causes multiple organ impairment (Stryjewski *et al.* 2003). However, detailed mechanism of its colonization remain unknown despite emerging clinical documentations on this respiratory pathogen (Atkinson *et al.* 2006; Daneshvar *et al.* 2001; Stryjewski *et al.* 2003). To date, *Pandoraea* sp. is recognized as one of the less studied CF pathogens that requires further investigations particularly in its bacterial pathogenicity (Callaghan & McClean 2012).To aggravate the situation, *Pandoraea* sp. was often misidentified in many clinical laboratories leading to the lack of clinical documentation on its virulence potential (Hogardt *et al.* 2009). On the other hand, *Pandoraea* sp. have considerable attractions in biotechnological applications with various degradation abilities such as lignin degradation (Shi *et al.* 2013), polychlorinated biphenyls (PCBs) biodegradation (Dhindwal *et al.* 2011), and sulphur oxidation (Anandham *et al.* 2008).

Previously, we reported the first documentation of *P. pnomenusa* RB38 isolated from a non-operating landfill site which produce C8-HSL (Ee *et al.* 2014b). As there is no report of AHL synthase in the *Pandoraea* genus, we sought to identify the presence of the AHL synthase in the genome of *P. pnomenusa* RB38 and further study it. We started the experiment by sequencing the complete genome of *P. pnomenusa* RB38 to provide the vital groundwork to understand this strain comprehensively prior to gene hunting. As QS is well-known to regulate various gene expressions such as virulence factors, identification of the LuxI/R homologs will be useful for further investigations on the QS-regulated gene expression. To our best knowledge, this is the first documentation of the QS system in the genus of *Pandoraea*.

## METHODS

### Bacterial Strains and Culture Conditions

LB medium (Scharlau, Spain) was used as the only culture media in the experiment. The AHLs biosensors used in this experiment was *Chromobacterium violaceum* CV026, *Escherichia coli* [pSB401] and *E. coli* [pSB 1142] while *Erwinia carotovora* GS101 and *E. carotovora* PNP22 was used as the positive and negative control for screening of AHLs production. All isolates were cultured routinely in Luria-Bertani (LB) agar or broth in 28°C with exception to *Escherichia coli* [pSB401], *E. coli* [pSB 1142] and *E. coli* BL21(DE3)pLysS, which were cultured aerobically at 37°C.

### Complete Genome Sequencing

Complete genome sequencing was performed using Pacific Biosciences (PacBio) RS II Single Molecule Real Time (SMRT) sequencing technology (Pacific Biosciences, Menlo Park, CA) as described previously (Chan, Yin & Lim 2014; Ee *et al.* 2014c). Briefly, the prepared 10-kb template library was sequenced on 4 single molecule real time (SMRT) cells using P4-C2 chemistry. *De novo* assembly was performed by filtering insert reads using RS_filter protocol (version 2.1.1) prior to assembly with Hierarchical Genome Assembly Process (HGAP) workflow in SMRT portal (version 2.1.1). Gene prediction was conducted using Prodigal version 2.60 (Hyatt *et al.* 2010).

### Whole Genome Mapping

Whole genome mapping was performed using OpGen Argus® system (OpGen, Gaithersburg, MD) according to manufacturer’s instructions. Briefly, high molecular weight DNA was isolated from single colony of sample strain using Argus High Molecular Weight (HMW) DNA Isolation Kit. DNA quality and concentration were determined using the Argus QCard kit. Single DNA molecules were then flowed through a microfluidic channel that was formed by Channel Forming Device (CFD), and were immobilized on charged glass surface. By using Enzyme Chooser software, BamHI was selected as the optimal restriction endonuclease for *P. pnomenusa* RB38, based on the FASTA-formatted sequence generated from PacBio RS II sequencing technology. The DNA molecules were digested on the glass surface to maintain the fragment order, and were stained with fluorescence dye. The image of DNA fragments were captured using fluorescence microscopy and fully automated image-acquisition software. The single-molecule maps were assembled by overlapping DNA fragment patterns to produce a whole genome map (WGM) with a minimum of 30X coverage. The WGM was aligned with PacBio FASTA-formatted sequences using sequence placement tool in MapSolver software (OpGen, Gaithersburg, MD).

### Identification of Putative *N*-acyl Homoserine Lactone Synthase and Gene Cloning

The predicted open reading frames (ORF) were further annotated by comparing against NCBI-NR (ftp://ftp.ncbi.nlm.nih.gov/blast/db/) and Uniprot databases (http://www.uniprot.org/) to locate the AHL synthase. The putative *ppnI* sequence was then sent to GenScript Inc. for gene cloning service where it was cloned in pUC57 vector (GeneScript, Piscataway, NJ) prior to direct cloning into pGS-21a expression vector. The resulting pGS-21a::*ppnI* plasmid was transformed into competent *E. coli* BL21(DE3)pLysS. Ampicillin (100μg/ml) and chloramphenicol (34μg/ml) (CalBioChem, Merck Millipore, Billerica, MA) were added for transformation selection purpose.

### Screening of AHL Production

Preliminary screening of AHL was performed by streaking transformed *E. coli* with gene of interest against *C. violaceum* CV026 biosensor prior to 37°C incubation overnight. *E. coli* harboring only vector pGS-21a without the gene of interest was included as negative control.

AHL extraction was performed as previously described (Ee *et al.* 2014a). Briefly, planktonic culture of transformed *E. coli* with gene of interest was extracted twice with equal volume of acidified ethyl acetate (0.1% glacial acetic acid) and left to complete desiccation until further analysis (Ortori *et al.* 2011).

AHL profile was confirmed using LC-MS/MS triple quadrupole mass spectrometry (Agilent 1290 Infinity LC and Agilent 6490 Triple Quadrupole LC/MS systems, Agilent Technologies, Santa Clara, CA) as described previously (Ee *et al.* 2014a; Lim *et al.* 2014). AHL detection was performed using precursor ion mode where the precursor ion *m/z* value was scanned from 80 to 400. Agilent MassHunter software was used for data analysis.

### Thin Layer Chromatography

Thin layer chromatography was conducted with loading of 25μL of extracted AHLs (in 100μL of ACN) on activated reverse phase C18 TLC plate (TLC aluminium sheets 20cm × 20cm, RP-18 F254s, Merck, Darmstadt, Germany) (Shaw *et al.* 1997). Synthetic AHLs of *N*-octanoyl-_L_-homoserine lactone (C8-HSL) (Sigma–Aldrich, St Louis, MO) were included as positive control and the chromatography was performed in 60% methanol: 40% water volume. Once completed, the TLC plate was air-dried and seeded with overnight culture of CV026 biosensor before it was left for overnight incubation (Chen *et al.* 2013; Lim *et al.* 2014).

## RESULTS AND DISCUSSION

### Complete Genome Sequencing

In this study, PacBio RSII SMRT sequencing technology was used as the sequencing platform in which the genome of *P. pnomenusa* RB38 was assembled into a single contig (GenBank accession number CP007506.1). With an average coverage of 190-fold, 4755 ORFs were revealed in the 5.3797Mb complete genome of *P. pnomenusa* RB38. By using Gepard (Krumsiek, Arnold & Rattei 2007), a dot matrix analysis was performed on the FASTA formatted sequence file of the genome which confirmed the circular topology of the assembly (data not shown).

This complete genome was then validated using OpGen WGM processed with BamHI. Optical mapping is commonly used as one of the laboratory techniques to provide a structural scaffold for contigs orientation as well as to visually identify errors in genome assemblies by using constructed whole genome restriction maps (Nagarajan, Read & Pop 2008). Perfect alignment of the WGM (5.146Mb) constructed with the complete genome assembly of *P. pnomenusa* RB38 confirmed the accuracy of the finished genome sequence.

### Multilocus Sequence Typing (MLST)

*Pandoraea* spp. belong to the beta-subclass of Proteobacteria with *Burkholderia* and *Ralstonia* as the closest neighbor (Coenye *et al.* 2000). In clinical microbiology laboratory, *Pandoraea* spp. is often misidentified as *Burkholderia cepacia* complex (Bcc) or *Ralstonia* spp or initially reported as non-fermentative Gram-negative bacilli (Aravena-Román 2008; Coenye *et al.* 2001). Initial annotation of *P. pnomenusa* RB38 complete genome using Rapid Annotation using Subsystem Technology (Version 4.0) (http://rast.nmpdr.org/rast.cgi) misidentified *Burkholderia* sp. CCGE1001 as the closest identity. However, isolate identification performed in previous study using 16S rDNA sequencing and Matrix-assisted Laser Desorption Ionization Time-of-Flight Mass Spectrometry (MALDI-TOF MS) identified strain RB38 as *P. pnomenusa* [24].

Multilocus sequence typing (MLST) is considered as the “gold standard” of accurate typing and identification of bacterial species (Larsen *et al.* 2012). As there is currently no MLST studies available for *Pandoraea* species, we employed the *Burkholderia* MLST database as a reference and also to study the possibility of employing *Burkholderia* MLST in distinguishing *Pandoraea* genus effectively. With the availability of the whole genome sequence data for strain RB38, Multilocus Sequence Typing (MLST) analysis was performed (**Table 1**), where seven conserved housekeeping genes (*atpD*, *gltB, gyrB, recA, phaC, lepA* and *trpB*) from genomic sequences of *P. pnomenusa* RB38 were blasted against NCBI-NR database for the nearest identity. As expected, all conserved housekeeping genes with exception of *lepA* gene successfully distinguished *Pandoraea* sp. as the closest organism [24].

**Table 1:** Multi-Locus Sequence Typing Analysis of *P. pnomenusa* RB38. Seven housekeeping genes in *P. pnomenusa* RB38 were analyzed where six out of seven conserved genes of *Pandoraea* were successfully distinguished from *Burkholderia.* All six MLST sequences shows an expect-value of 0.0, which reflects high similarity except for acetoacetyl-CoA reductase (*phaC*) that gives the value of expect-value169 for *Pandoraea sp*. SD6-2 and expect-value130 for *Burkholderia phymatum* STM815. As shown below, *P. pnomenusa* RB38 is highly similar to *Pandoraea sp*. SD6-2 with higher identities.

| Genes | First Match |  |  | Second Match |  |  |
| --- | --- | --- | --- | --- | --- | --- |
|  | Strain | Identities | Expect | Strain | Identities | Expect |
| **ATP synthase beta chain (*atpD*)** | *Pandoraea* sp. SD6-2 | 444/463 (96%) | 0 | *Burkholderia vietnamiensis* G4 | 425/464 (92%), | 0 |
| **Glutamate synthase large** | *Pandoraea* sp. SD6-2 | 1496/1563 (96%), | 0 | *Burkholderia terrae* | 1308/1566 (84%) | 0 |

**Table 1: Multi-Locus Sequence Typing Analysis of *P.**
| subunit (*gltB*) |  |  |  |  |  |  |
| --- | --- | --- | --- | --- | --- | --- |
| **DNA gyrase subunit B(*gyrB*)** | *Pandoraea* sp. SD6-2 | 744/825 (90%), | 0 | *Burkholderia ambifaria* MC40-6 | 657/830 (79%) | 0 |
| **Recombinase A (*recA*)** | *Pandoraea* sp. SD6-2 | 338/349 (97%) | 0 | *Burkholderia rhizoxinica* HKI 454 | 305/342 (89%) | 0 |
| **Acetoacetyl-CoA reductase(*phaC*)** | *Pandoraea* sp. SD6-2 | 224/247 (91%) | e-169 | *Burkholderia phymatum* STM815 | 173/247 (70%) | e-130 |
| **GTP binding protein (*lepA*)** | Not Found | Not Found | Not Found | Not Found | Not Found | Not Found |
| **Tryptophan synthase subunit beta (*trpB*)** | *Pandoraea* sp. SD6-2 | 384/400 (96%) | 0 | *Ralstonia pickettii* 12J | 335/393 (85%), | 0 |

### Identification and *in silico* Analysis of *luxI/R*-Type QS Genes

We previously reported the QS activity of *P. pnomenusa* RB38 (Ee *et al.* 2014b). In this study, we identified the putative *luxI* and *luxR* homologs from the annotated genome. Firstly, a 786 bp putative *N-*acyl homoserine lactone synthase (DA70_23485) (designated as *ppnI* gene) was identified. Conserved domain analysis of the predicted proteome of this gene indicated presence of autoinducer synthase domain (PFAM signature: PF00765) which further confirmed that this gene is a genuine LuxI homolog. Further, a 702 bp putative cognate LuxR homolog (DA70_23490) (designated as *ppnR* gene) located in close proximity and in a convergent transcriptional orientation to the *ppnI* gene was also manually identified (**Figure 1)**. Presence of LuxR homolog in close proximity to the LuxI homolog is commonly observed in the typical LuxI/LuxR-type QS circuit (Schaefer *et al.* 2013). In order to confirm the authenticity of this putative LuxR homolog, the predicted protein sequence was scanned and was confirmed to contain the universal conserved domain organization of LuxR proteins namely: the autoinducer binding domain (PFAM03472) and C-terminal DNA-binding domain of LuxR-like proteins (cd06170) (Choi & Greenberg 1992; Fuqua, Parsek & Greenberg 2001; Hanzelka & Greenberg 1995).

**Figure 1.**
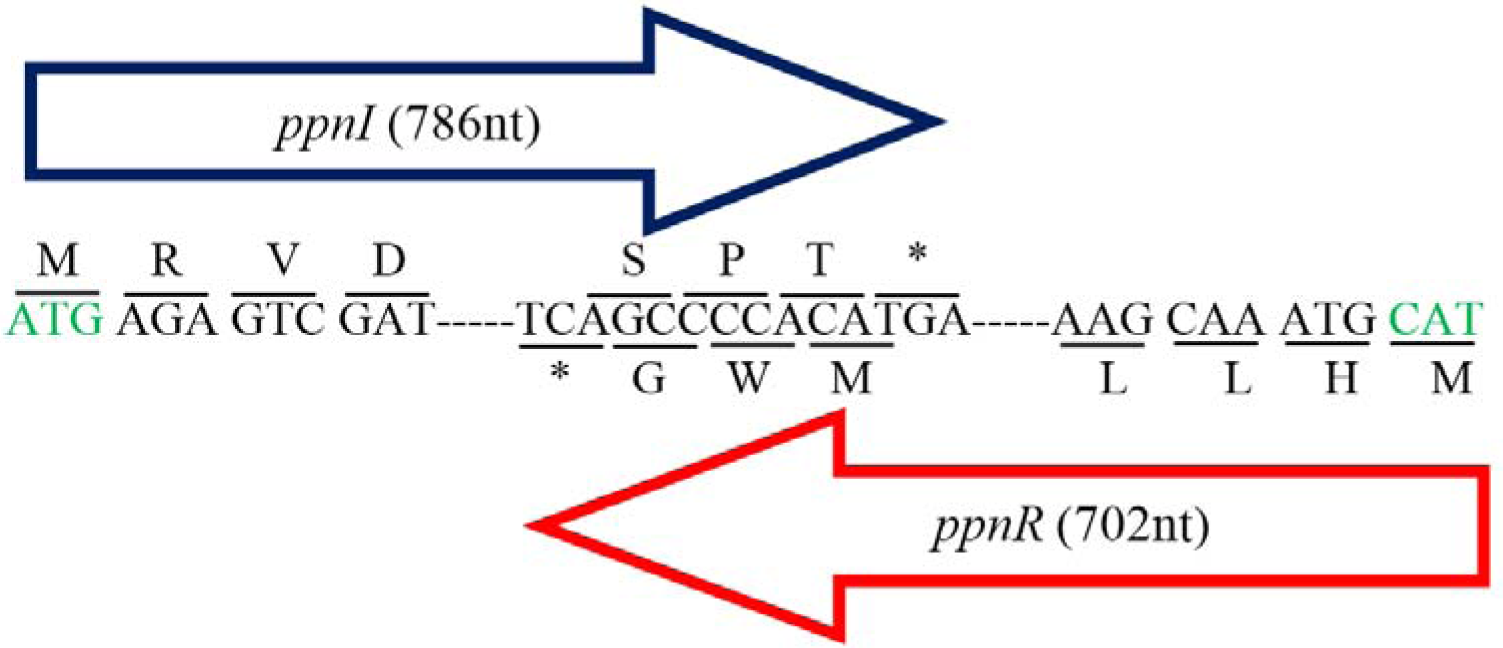
Gene map showing organization of *ppnR* (*luxR* homolog) and *ppnI* (*luxI* homolog). The direction of the arrows indicated the orientation of both genes where *ppnI* is in 5’-3’ direction while *ppnR* is in 3’-5’ direction. A line was used to indicate the nucleotide sequences and its respective amino acid sequence. Start codon, Methionine (M) was represented by green font; while asterisk represented stop codon (TGA). The *ppnR* and *ppnI* genes sequences had been deposited in GenBank database with GenBank accession number AHN77102.1 and AHN77101.1, respectively.

In addition, further search in the genome also indicated presence of an additional putative *luxR* homologous gene (designated as *ppnR*2) which was not associated with a *luxI* homolog and is therefore referred in this study as a putative orphan LuxR regulator. Orphan LuxR is hypothesized to occur as a result of genes re-organizations, horizontal gene transfer or independent evolution of transcriptional regulatory circuits (Patankar & González 2009b). Various studies have reported identification of orphan LuxR in numerous bacteria and orphan LuxR was also found to interact with AHLs to regulate a variety of gene expression (Malott *et al.* 2009; Patankar & González 2009a; Subramoni & Venturi 2009)

Furthermore, phylogenetic analyses based on amino acid sequences performed indicated that both the PpnI/PpnR1 pair and the orphan PpnR2 are distant from LuxI or LuxR homologues of its closest phylogenetic neighbour, both *Burkholderia* and *Ralstonia* species (**Figures 1, 2, 3**). To the best of our knowledge, this is the first documentation of LuxI/R homologs of the *Pandoraea* species.

**Figure 2.**
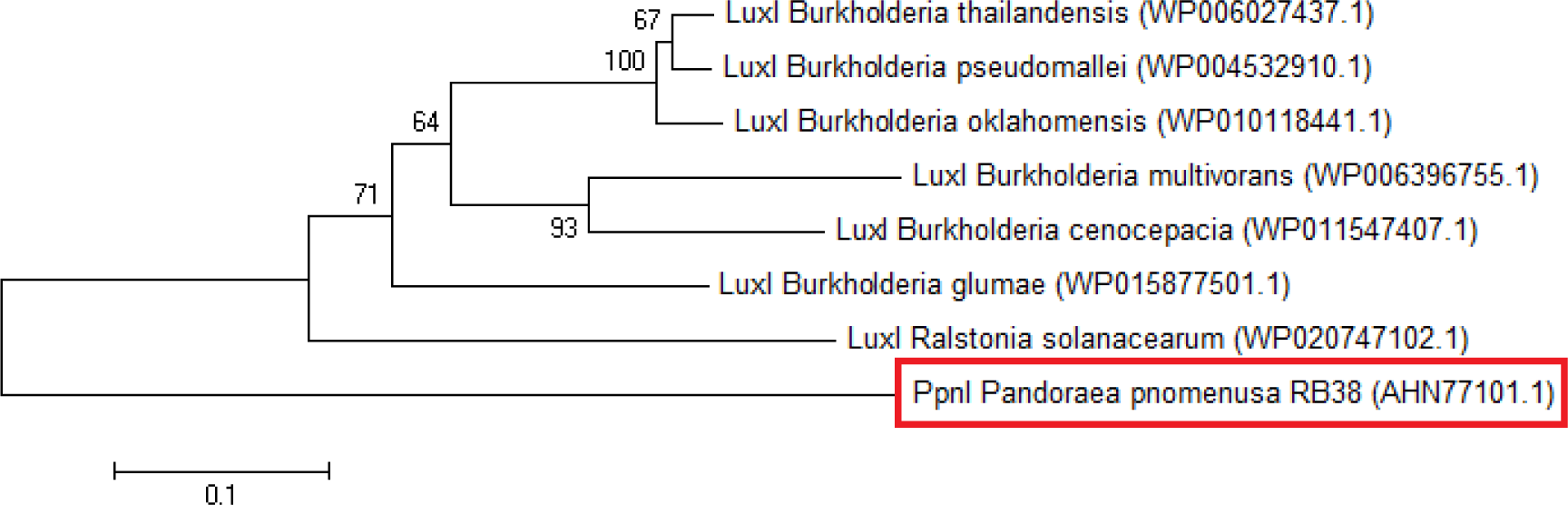
Phylogenetic tree of PpnI. Neighbor-Joining method (Saitou & Nei 1987) was used in MEGA6 (Tamura *et al.* 2011) where bootstrap tests (1000 replicates) were shown next to the branches (Felsenstein 1985). This analysis involved 8 amino acid sequences with their GenBank accession numbers as listed: LuxI *Burkholderia thailandensis* (WP006027437.1), LuxI *Burkholderia pseudomallei* (WP004532910.1), LuxI *Burkholderia oklahomensis* (WP010118441.1), LuxI *Burkholderia multivorans* (WP006396755.1), LuxI *Burkholderia cenocepacia* (WP015877501.1), LuxI *Burkholderia glumae* (WP015877501.1), LuxI Ralstonia solanacearum (WP020747102.1) and PpnI *Pandoraea pnomenusa* RB38 (AHN77101.1).

**Figure 3.**
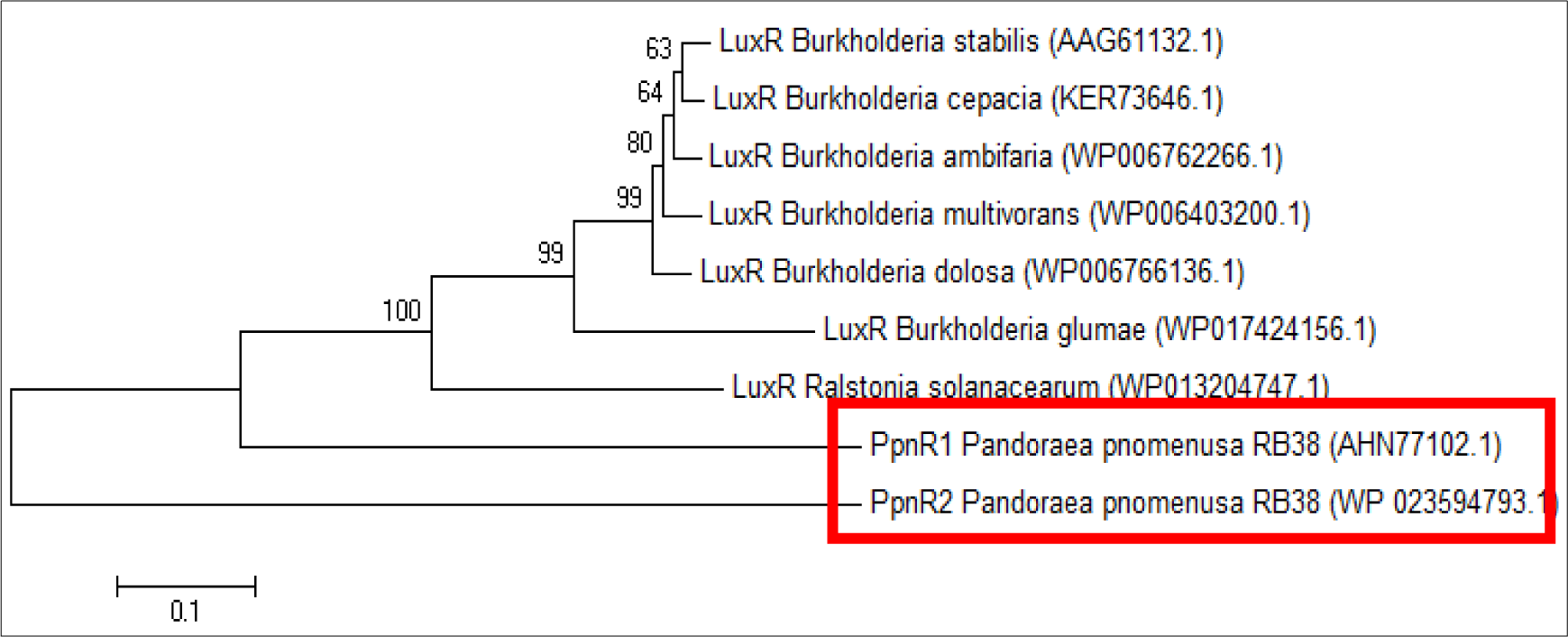
Phylogenetic tree of PpnR. Neighbor-Joining method (Saitou & Nei 1987) was used in MEGA5 (Tamura *et al.* 2011) where bootstrap tests (1000 replicates) were shown next to the branches (Felsenstein 1985). This analysis involved 9 amino acid sequences with their GenBank accession numbers as listed: LuxR *Burkholderia stabilis* (AAG61132.1), LuxR *Burkholderia cepacia* (KER73646.1), LuxR *Burkholderia ambifaria* (WP006762266.1), LuxR *Burkholderia multivorans* (WP006403200.1), LuxR *Burkholderia dolosa* (WP006766136.1), LuxR *Burkholderia glumae* (WP017424156.1), LuxR *Ralstonia solanacearum* (WP013204747.1), PpnR1 *Pandoraea pnomenusa* RB38 (AHN77102.1) and PpnR2 *Pandoraea pnomenusa* RB38 (WP023594793.1).

### Functional Study of Putative *ppnI* Gene

For functional studies, we cloned the putative *ppnI* into a pGS-21a expression vector and subsequently transformed the pGS-21a::*ppnI* plasmid into competent *E. coli* BL21(DE3)pLysS. AHL screening were performed using *C. violaeum* CV026 biosensor with *E. coli* BL21(DE3)pLysS::*ppnI.* The result of the cross-streak bioassay demonstrated activation of purple violacein secretion of *C. violaeum* CV026 (**Figure 4A**) as well as chemiluminescence activity of *E. coli* [pSB 401] indicating the production of short chain AHLs by the *ppnI* gene (**Figure 4B**). Besides that, formation of a sole purple violacein spot on CV026 lawn which correspond to the same retention time of the synthetic C8-HSL suggested that the *ppnI* is responsible for the production of C8-HSL in *P. pnomenusa* RB38 (**Figure 5**). The AHL profile of *ppnI* was further verified using LC/MS mass spectrometry system and only C8-HSL was detected in the supernatant of transformed *E.coli* BL21 suggesting that *ppnI* is indeed the functional LuxI synthase of *P. pnomenusa* RB38.

**Figure 4.**
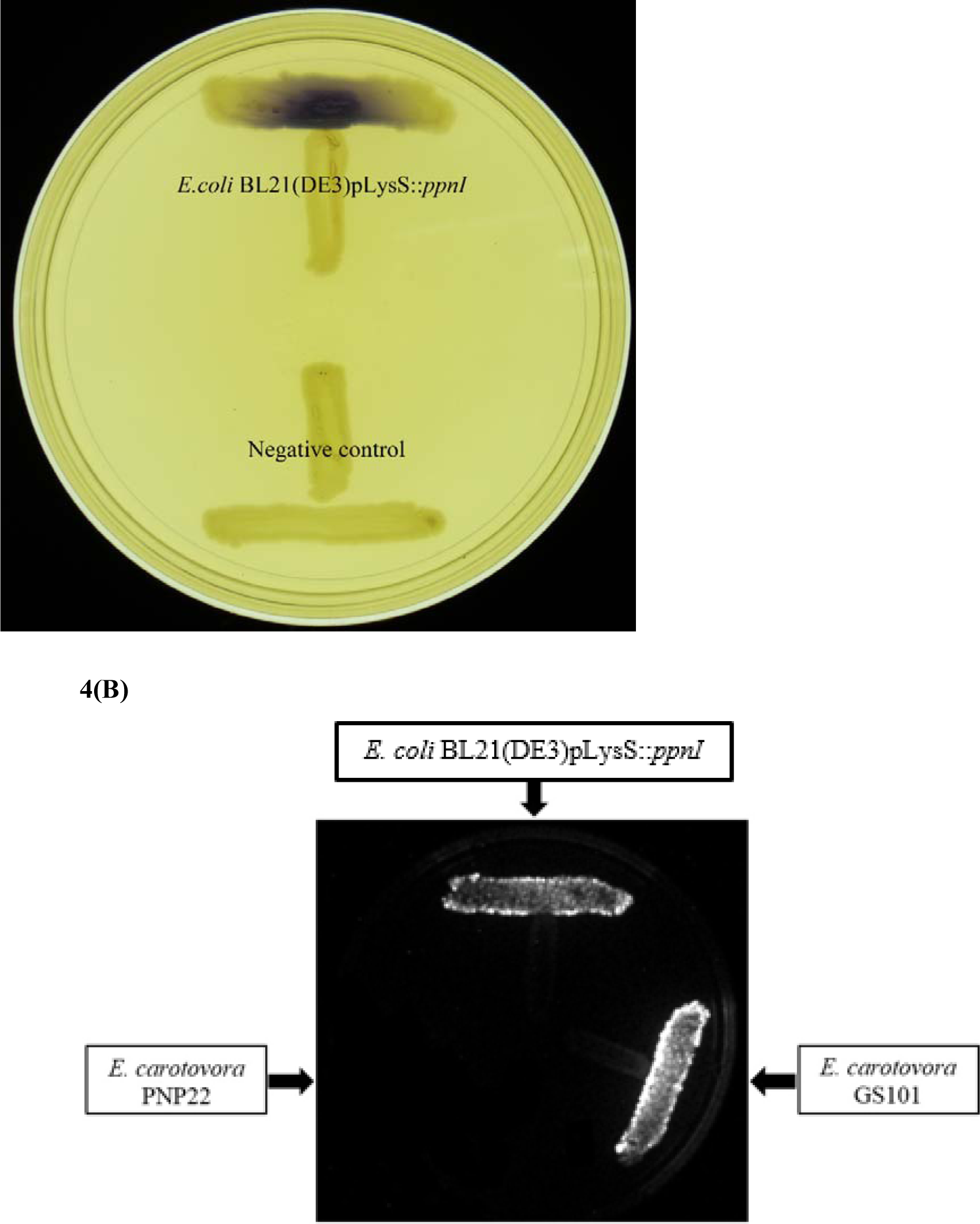
Cross Streaking bioassay. **(A) CV026 bioassay.** Purple pigmentation indicated secretion of short chain AHLs from *E.coli* BL21(DE3)pLysS::*ppnI*. **(B) *E. coli* [pSB 401] chemilumiscence bioassay.** Expression of chemiluminescence activity in *E. coli* [pSB 401] demonstrated the detection of short chain AHLs. *E. carotovora* GS101 and *E. carotovora* PNP22 served as the positive and negative control respectively in both experiment.

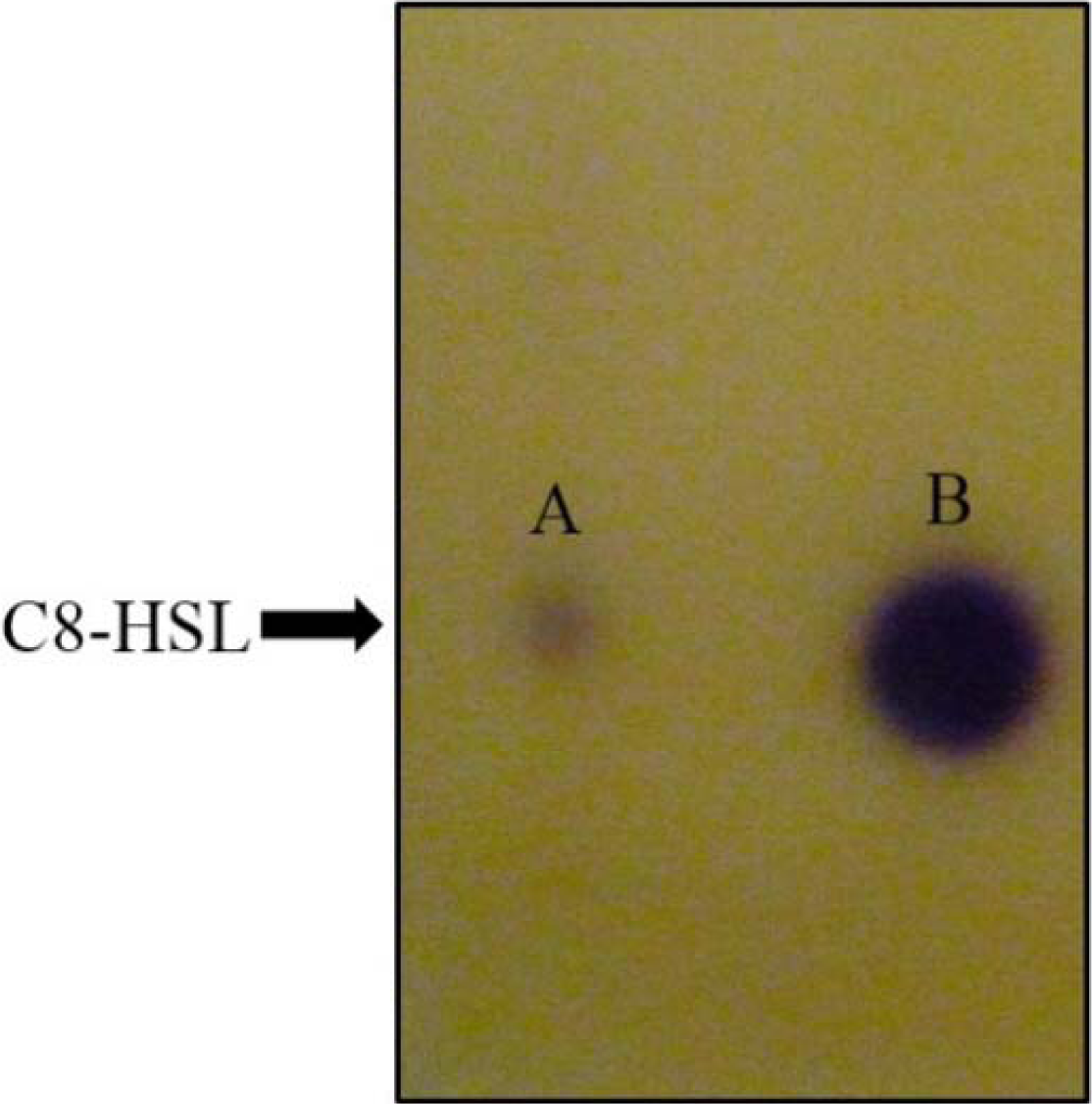

## Conclusion

Here, we reported the complete genome of *P. pnomenusa* RB38 and the discovery of its AHL synthase, designated as *ppnI* gene and its LuxR homolog receptor, *ppnR* gene, as well as an additional orphan LuxR regulator, *ppnR*2 gene. Short chain AHL, C8-HSLwas detected in the spent culture supernatant of *E.coli* BL21(DE3)pLysS::*ppnI* which confirmed that *ppnI* gene is a functional AHL synthase.

## ADDITIONAL INFORMATION AND DECLARATIONS

### Author Contributions

Yan-Lue Lim, Robson Ee, Kah-Yan How, Siew-Kim Lee and Wai-Fong Yin performed the research, data analysis and prepared for manuscript. Kok-Gan Chan designed, supervised and approved the experiments.

